# A family of cell wall transglutaminases is essential for appressorium development and pathogenicity in *Phytophthora infestans*

**DOI:** 10.1101/2021.11.23.469665

**Authors:** Maja Brus-Szkalej, Christian B. Andersen, Ramesh R. Vetukuri, Laura J. Grenville-Briggs

## Abstract

Transglutaminases (TGases) are enzymes highly conserved among prokaryotic and eukaryotic organisms, where their role is to catalyse protein cross-linking. One of the putative TGases of *Phytophthora infestans* has previously been shown to be localised to the cell wall. Based on sequence similarity we were able to identify six more genes annotated as putative TGases and show that these seven genes group together in phylogenetic analysis. All of the seven proteins are predicted to contain transmembrane helices and both a TGase domain and a MANSC domain, the latter of which was previously shown to play a role in protein stability. Chemical inhibition of transglutaminase activity and silencing of the entire family of the putative cell wall TGases are both lethal to *P. infestans* indicating the importance of these proteins in cell wall formation and stability. The intermediate phenotype obtained with lower drug concentrations and less efficient silencing displays a number of deformations to germ tubes and appressoria. Both chemically treated and silenced lines show lower pathogenicity than the wild type in leaf infection assays. Finally, we show that appressoria of *P. infestans* possess the ability to build up turgor pressure and that this ability is decreased by chemical inhibition of TGases.

## Introduction

Transglutaminases (TGases) are a group of enzymes (EC 2.3.2.13) that catalyse acyl transfer between γ-carboxamide groups of peptide-bound glutamine and a wide variety of acceptor substrates, usually primary amino acids (Folk, 1980). Of most interest and hence the best studied reaction catalysed by TGases is the interaction with ε-amino groups of peptide-bound lysine residues resulting in inter- or intramolecular protein cross-linking (ε-(γ-glutamyl) lysine cross-link) (Folk, 1980, Lorand and Graham, 2003). Although first discovered and extracted from animal tissue (Sarkar et al., 1957, Folk and Cole, 1966), TGases are widely conserved among both prokaryotic and eukaryotic organisms (Martins et al., 2014, Icekson and Apelbaum, 1987). The conservation of enzymatic properties in different organisms has therefore allowed for replacement of the previously used guinea pig TGase by the cheaper and ethically preferable microbial transglutaminase (MTG) in food manufacturing (Martins et al., 2014, Kieliszek and Misiewicz, 2014). The biological roles of transglutaminases are still best characterised for humans, where all nine TGases have been assigned to different organs and a specific function has been assigned to eight of them (Eckert et al., 2014). The microbial TGases identified to date do not share a significant sequence similarity to any of the other characterised TGase classes. Their biological functions remain largely unknown and they are essentially unstudied beyond their early identification (Makarova et al., 1999, Giordano and Facchiano, 2019). An *in silico* study by Makrova et al. (1999) identified a class of microbial proteins, with only one characterised member – a single protease, that shares a significant sequence similarity with animal TGases. They thus propose proteases as a common ancestor of all transglutaminases. Interestingly, although plant TGases do not show high sequence similarity to those from animals, they do resemble them structurally and can interact with animal TGase antibodies (Beninati et al., 2013). The main functions of plant TGases are usually linked to photosynthesis, responses to stress, senescence and programmed cell death and it is believed they do so through stabilisation and modification of various proteins involved in these processes (Zhong et al., 2019, Serafini-Fracassini and Del Duca, 2008).

The first oomycete TGase to be characterised was found thanks to its role in the induction of immune responses in plants. A 42kDa cell wall glycoprotein (GP42) of *Phytophthora sojae*, previously shown to contain thirteen amino acid peptide - Pep13 that acts as an elicitor of defence mechanisms in parsley (Nürnberger et al., 1994, Hahlbrock et al., 1995), was characterised as a Ca^2+^- dependent TGase and its homologs were found in all *Phytophthora* species, including *P. infestans* (Brunner et al., 2002). Although the GP42 protein showed enzymatic and biochemical similarities to human transglutaminases its folding patterns were shown to be novel and to not resemble any characterised proteins (Reiss et al., 2011).

We have previously shown that peptides from the putative TGase encoded by the *P. infestans* gene PITG_22117 are found predominantly in the cell wall of germinated cysts and appressoria (Grenville-Briggs et al., 2010) and that the gene is highly expressed in cysts, appressoria and during the early stages of infection of potato plants (Grenville-Briggs et al., 2010, Resjö et al., 2017). These results suggest a role for TGases in the development of infectious structures and hence the pathogenicity of *P. infestans*. To test this hypothesis, in the current study we have performed an *in silico* analysis of the family of putative TGases with the highest sequence similarity to PITG_22117. We have identified six additional genes and transiently silenced all seven of them to screen for possible developmental changes due to the lack of expression of putative TGases. Silencing resulted in structural changes in appressorium formation which correlated with a reduction in pathogenicity. Similar phenotypes were observed upon treatment of cysts of *P. infestans* with the transglutaminase inhibitor cystamine.

## Results and Discussion

### There are 21 putative transglutaminases identified in Phytophthora infestans

In previous studies we have shown that at least one of the *P. infestans* transglutaminases is found in the cell wall of germinating cysts and appressoria (Grenville-Briggs et al., 2010). We have verified that the PITG_22117 gene encoding this protein is highly expressed in cysts and appressoria and at early infection stages – 6 hpi and 12 hpi, when the appressoria are formed on the leaf surface and start to penetrate the host cells (Resjö et al., 2017). Here, we have used an *in silico* approach to look for other TGases using sequence similarity searches. A BLASTn search yielded five additional genes, two of which belong to the M81 gene family: PITG_16956, PITG_16958, PITG_16959 (M81D), PITG_16963 (M81C), PITG_22104; and one gene labelled just as M81E. The M81 gene family was characterised as a family of elicitor-like proteins with divergent structures (Fabritius and Judelson, 2003). We also used string searches, such as “*Phytophthora infestans* transglutaminase” using the NCBI non-redundant (nr) Gene database and found 21 results with varying degrees of annotation. We have retrieved protein sequences of all of these 21 genes and constructed a maximum likelihood phylogenetic tree (Figure 1). The phylogenetic analysis of the protein sequences confirmed that proteins encoded by the genes found by BLASTn group together. From the phylogenetic analysis we have also identified two additional genes of high sequence similarity PITG_16953 and PITG_08335. Moreover, with the exception of PITG_22104, which is a short sequence with 100% similarity to PITG_16959 and thus a possible pseudogene, all of the six genes with high similarity to PITG_22117 include the conserved sequence encoding the Pep13 peptide, an elicitor of plant defence responses (Nürnberger et al., 1994, Brunner et al., 2002). The PITG_22104 gene was excluded from further analysis. These findings are to some extent consistent with the study showing the M81 gene family to encode elicitors of which several are transglutaminases containing Pep13 (Fabritius and Judelson, 2003). While the M81 protein was reported to be mating specific, other members of the family are expressed at different life cycle stages. A more recent study from the same group reports PITG_13497 as induced over 100-fold during mating, where it is believed to play an important role in the synthesis of the very thick oospore cell wall (Niu et al., 2018).

**Figure 1.**
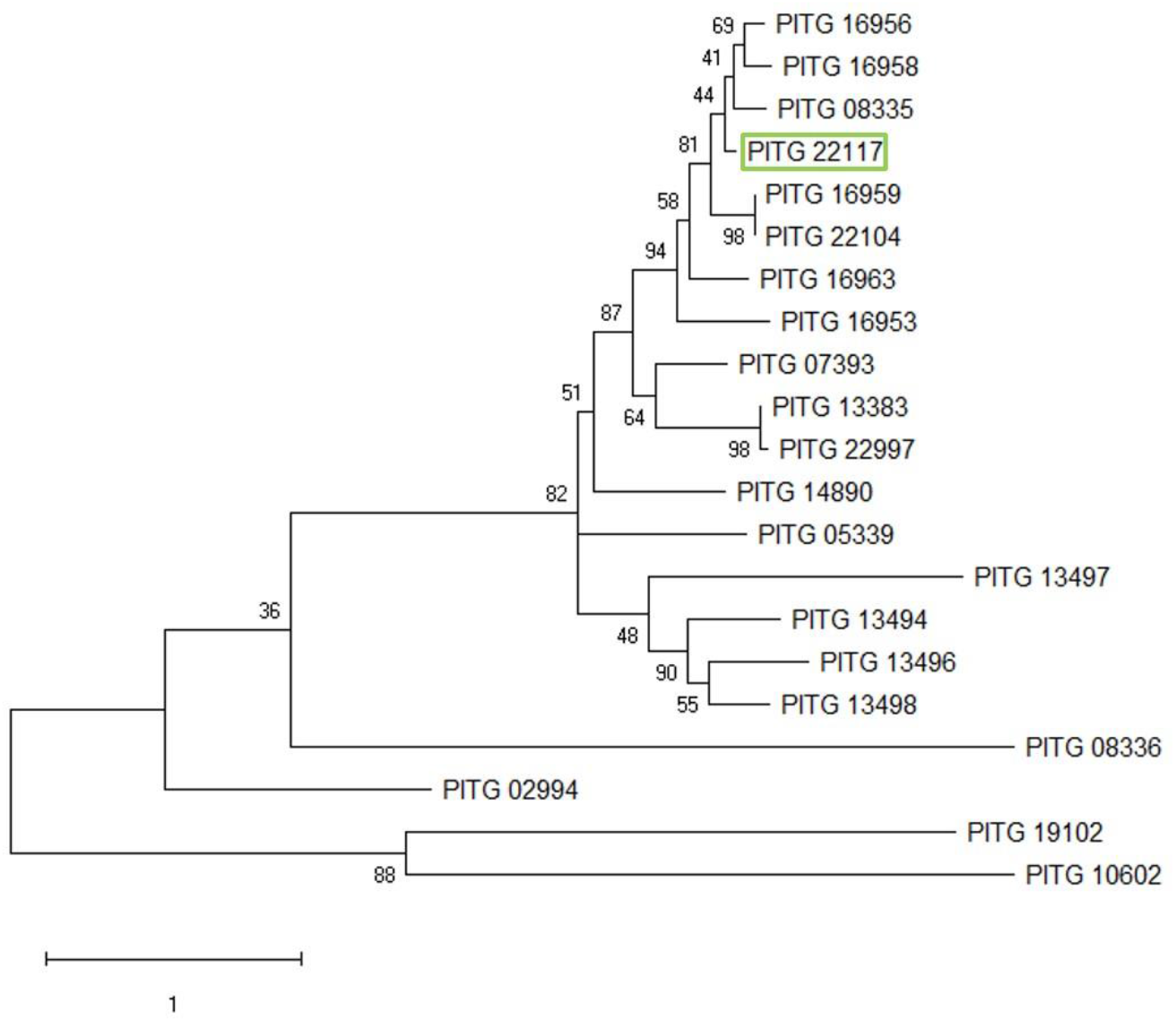
Maximum likelihood phylogenetic tree of *P. infestans* TGase proteins. Bootstrap, 1000 replicates. The scale bar represents number of substitutions per site.

### All Pep13 TGases are transmembrane elicitor proteins

The seven TGases containing the Pep13 peptide were functionally annotated *in silico*. The presence of a TGase elicitor domain (IPR032048) was predicted in all of them. Additionally, in all seven proteins the TGase domain overlapped largely with a MANSC – Motif At N terminus with Seven Cysteines domain (PTHR16021; Figure 2). All animal proteins containing the MANSC domain that have been reported so far contain both signal peptide and transmembrane helix regions, and the seven cysteines are suggested to form disulphide bonds that play a role in protein structure and stability (Guo et al., 2004). The presence of transmembrane helices was predicted by the use of transmembrane topology prediction software in proteins encoded by genes PITG_16956, PITG_16958, PITG_16959 and PITG_16953. All these proteins were also predicted to contain signal peptides at their N-termini. The lack of transmembrane helix or signal peptide prediction for the remaining proteins suggests that they may be entirely embedded in the cell wall, or that they may be cytoplasmic, or localised via non-classical secretion pathways. In the two shortest *P. infestans* proteins – PITG_22117 and PITG_08335 the TGase and MANSC domains cover the entire length of their sequences. PITG_16963 contains a stretch of 185 amino acids at its N-terminus for which there were no domains predicted, while the entire length of the remaining four proteins is predicted as a non-cytoplasmic domain (Figure 2), suggesting they may be localised to the cell wall via non-classical secretion. These predictions show that all seven members of this family are likely to be transmembrane proteins with putative TGase elicitor functions.

**Figure 2.**
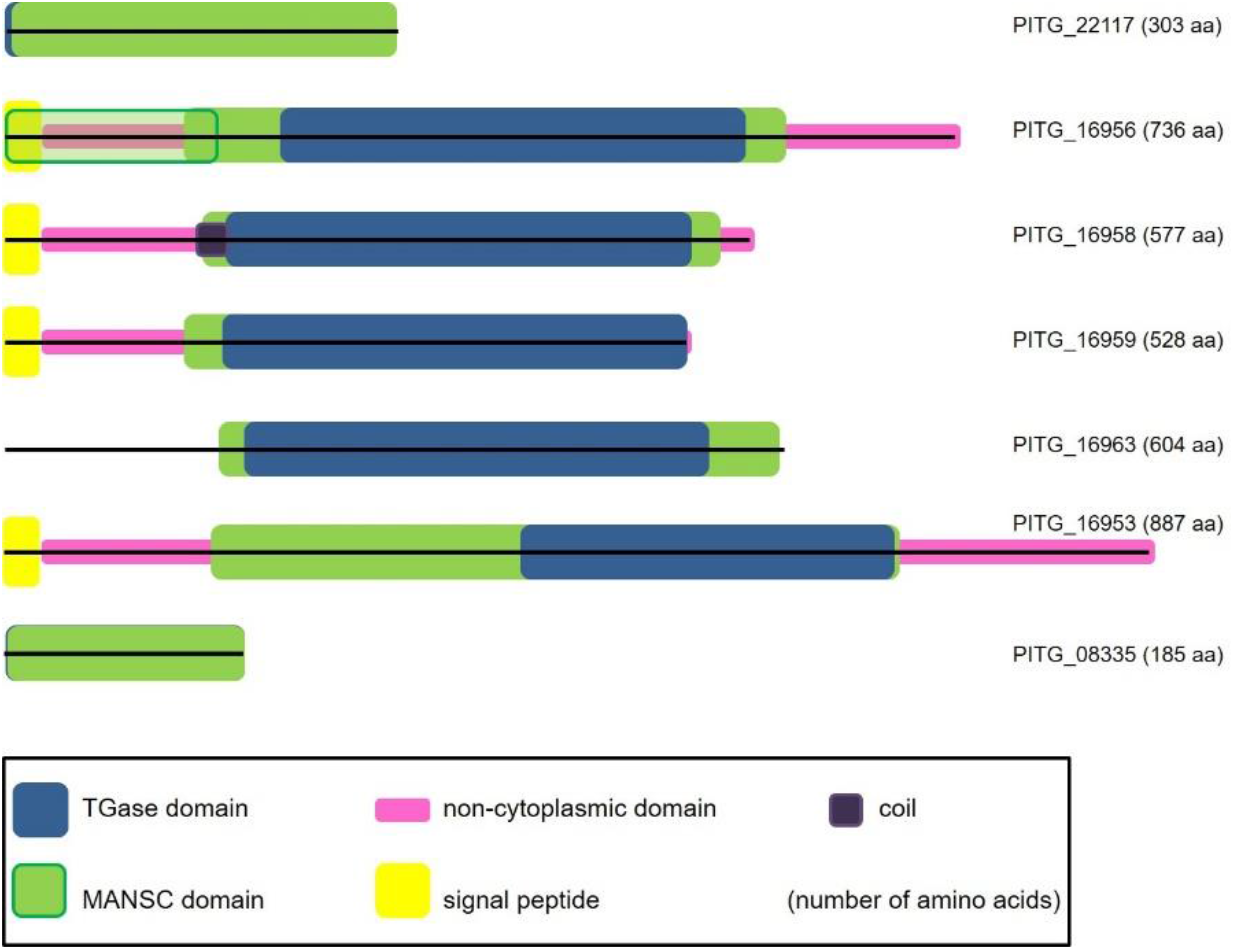
Functional domains in elicitor-TGases. The sequences are aligned at amino acid 1, to show the difference in length. Functional domains predicted by InterProScan are represented by blocks of different colours and drawn to scale (see legend at the bottom of the figure).

### The elicitor-TGases are highly expressed at early infection time points

The expression of all seven elicitor-TGase genes was compared in the asexual pre-infection stages (produced *in vitro*) and early infection time points (detached leaf assays) sampled at 6 hpi, 12 hpi and 24 hpi (Figure 3). Expression of PITG_16958 was undetectable in either *in vitro* or infection assays, which suggests that it might be expressed specifically in the sexual reproduction cycle, or at only very low levels. A recent study of *P. infestans* sexual reproduction revealed that gene expression could be strain-dependent and the level of expression in the oospores can vary for different strain combinations, with PITG_16958 highly upregulated in some of the oospores and downregulated in others (Tzelepis et al., 2020; and personal communication with the authors). These data also suggests that perhaps the PITG_16958 gene is not expressed in the 88069 strain that we used in the current study.

**Figure 3.**
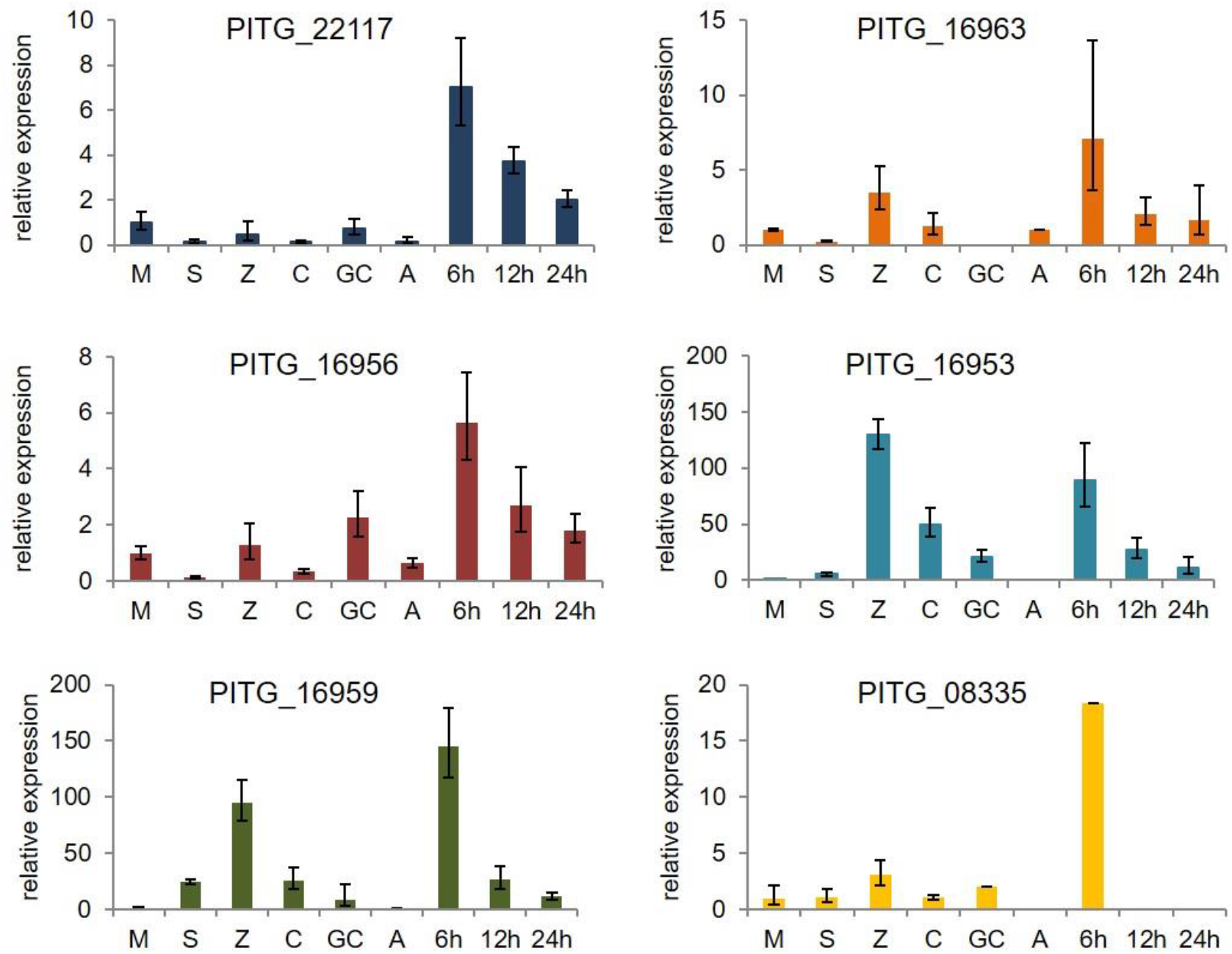
Expression profiles of the elicitor-TGases. The expression was calculated relative to ActA and calibrated to mycelium sample for which the expression was set to 1. M - mycelium, S – sporangia, Z – zoospores, C – cysts, GC – germinated cysts, A – appressoria, 6h, 12h, 24h – hours post inoculation with cyst solution. Gene PITG_16958 is not shown as ther was no signal detected for this gene. Error bars represent errors calculated using the modified Delta-Delta Ct method.

All of the other six genes showed the highest expression at 6 hpi on detached potato leaves, i.e. at the time point when the majority of appressoria are being formed on the leaf surface; and at 12 hpi when first penetration of the host cells occurs. PITG_16953 was the only gene in which the expression was the highest in the zoospores. The much higher level of gene expression at 6 hpi than in pre-infection stages in the other genes indicates that the expression of the genes is not only life cycle specific, but also induced by the presence of the host. PITG_22117 was the only gene in which no significant differences were found between the different pre-infection life cycle stages and the expression was induced only by the infection. Genes PITG_16959, PITG_16963 and PITG_16953 are most likely zoospore specific, since their expression was significantly higher at this stage than in any other part of the life-cycle tested. PITG_08335 exhibited higher expression in the zoospores, but the induction was only slight and much lower than in the other three genes (Figure 3). What is more, there was no signal read for this gene in appressoria, or at 12 hpi and 24 hpi indicating an absolute lack of expression during these stages (verified in three independent biological replicates). Since zoospores lack a cell wall and it is at the transition between the zoospore and cyst when the cell wall needs to be quickly formed *de novo* (Grenville-Briggs et al., 2008), the high level of gene expression points toward a role for these genes in cyst cell wall formation and possibly in the development of the (pre)infectious structures. PITG_16956 showed a more specific expression pattern being highly induced in the germinated cysts when the appressoria are formed. Nonetheless, the level of induction is generally much lower in this gene than in the zoospore specific ones (Figure 3). Therefore, we hypothesise that much higher levels of transglutaminases are necessary for the encystment and germination of cysts than for the formation of appressoria and thus several TGases are very highly expressed in the zoospores, whilst a lower induction is sufficient for the formation of appressoria.

### Chemical inhibition of TGases affects the growth and germination of P. infestans

Cystamine was previously reported to inhibit the activity of TGases in humans (Lorand and Graham, 2003, Jeitner et al., 2018) and fungi (Ruiz-Herrera et al., 1995, Iranzo et al., 2002). Cystamine treatment of fungi resulted in growth inhibition, morphological changes, influenced incorporation of peptides into the cell wall and inhibited the yeast-to-mycelium transition in dimorphic fungi (Reyna-Beltrán et al., 2019). However, the effects of cystamine on oomycete growth or development have not been tested before. We grew *P. infestans* in both liquid and solid media at a range of cystamine concentrations and were able to show that independent of the type of medium, 5 mM cystamine inhibits mycelial growth by about 40 %, 7.5 mM by 60%, 10 mM by 90 % and at 20 mM and higher concentrations there was no growth observed at all. Sporangial germination was also affected by the chemical treatment; 7.5 mM cystamine inhibited germination completely, while at 5 mM cystamine there were only single germ tubes found. The germ tubes at 2.5 mM and 1 mM cystamine had more deformations than the ones seen at lower concentrations (Figure 4).

**Figure 4.**
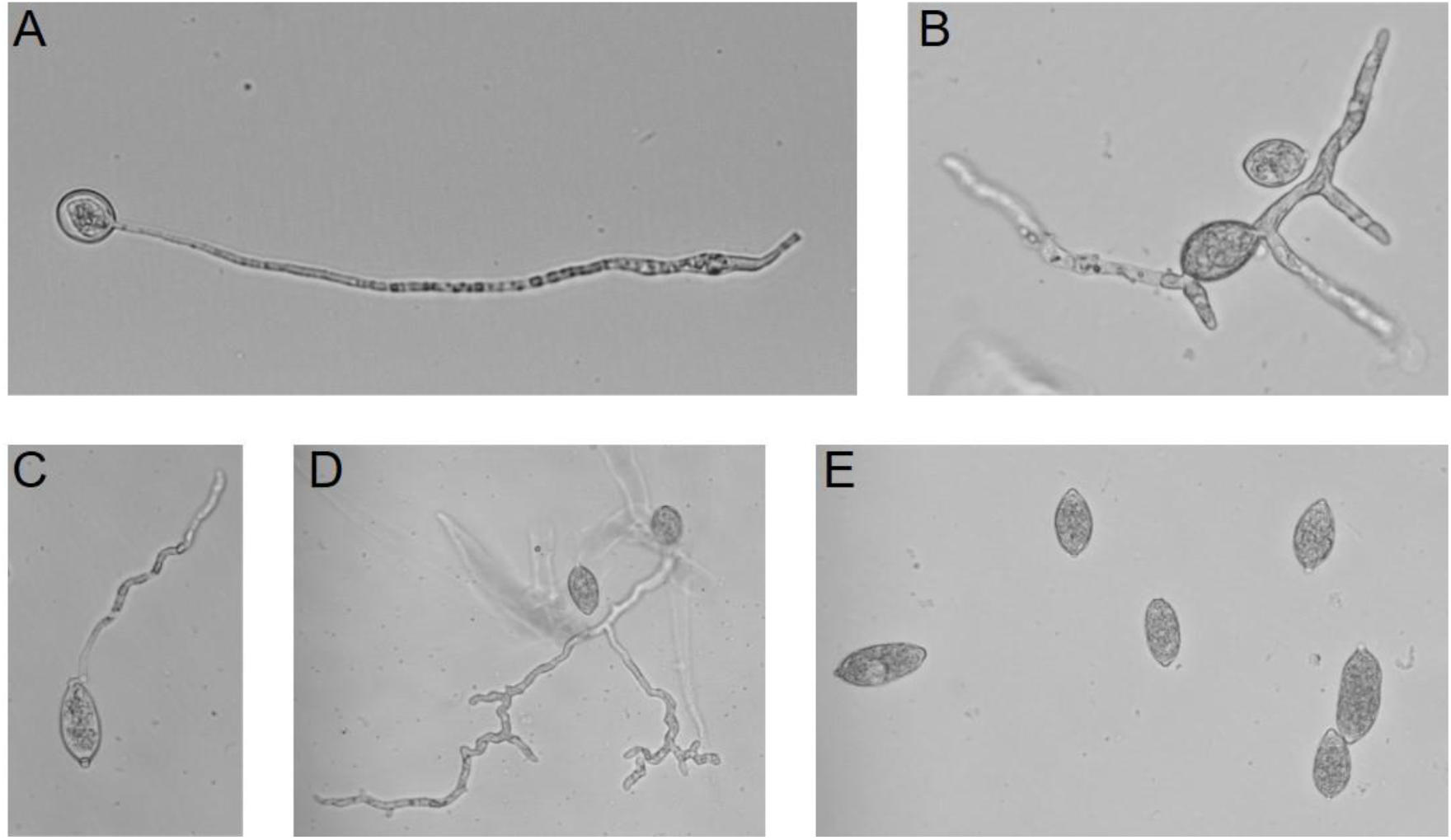
Effect of cystamine on sporangia germination. **A:** untreated sample, **B:** 1 mM cystamine, **C:** 2.5 mM cystamine, **D:** 5 mM cystamine, **E:** 7.5 mM cystamine. Panel **A** shows a healthy germinating sporangium. Addition of cystamine to the growth medium resulted in deformations of the germ tubes (**B, C, D**), lower percentage of germination (**D**) and at 7.5 mM (**E**) complete inhibition of germination.

To test the effect of cystamine on zoospore release, solid cultures were flooded with cystamine solutions of concentrations ranging between 0.5 and 200 mM. None of the concentrations had any effect on the release or motility of the zoospores. However, cysts produced from cystamine treated zoospores had a lower germination rate than untreated ones (Figure 5). At 0.5 mM the germination rate was about 50 % lower, while at 2.5 mM it decreased by 75 %. Interestingly, there were no differences in the rate of germination observed between 2.5 mM, 5 mM, 7.5 mM and 10 mM (Figure 5A). This suggests that germination may be only partially dependent on TGases, or that not all *P. infestans* TGases are sensitive to this inhibitor. Since most of the TGases appear to be cell wall localised, the hypothesis that the ones without the signal peptide are entirely embedded in the cell wall structure may somewhat explain the latter hypothesis, as they may not be accessible to the drug due to their location. An alternative might be that they are able to form complexes that protect their active sites from inhibition by this drug.

**Figure 5.**
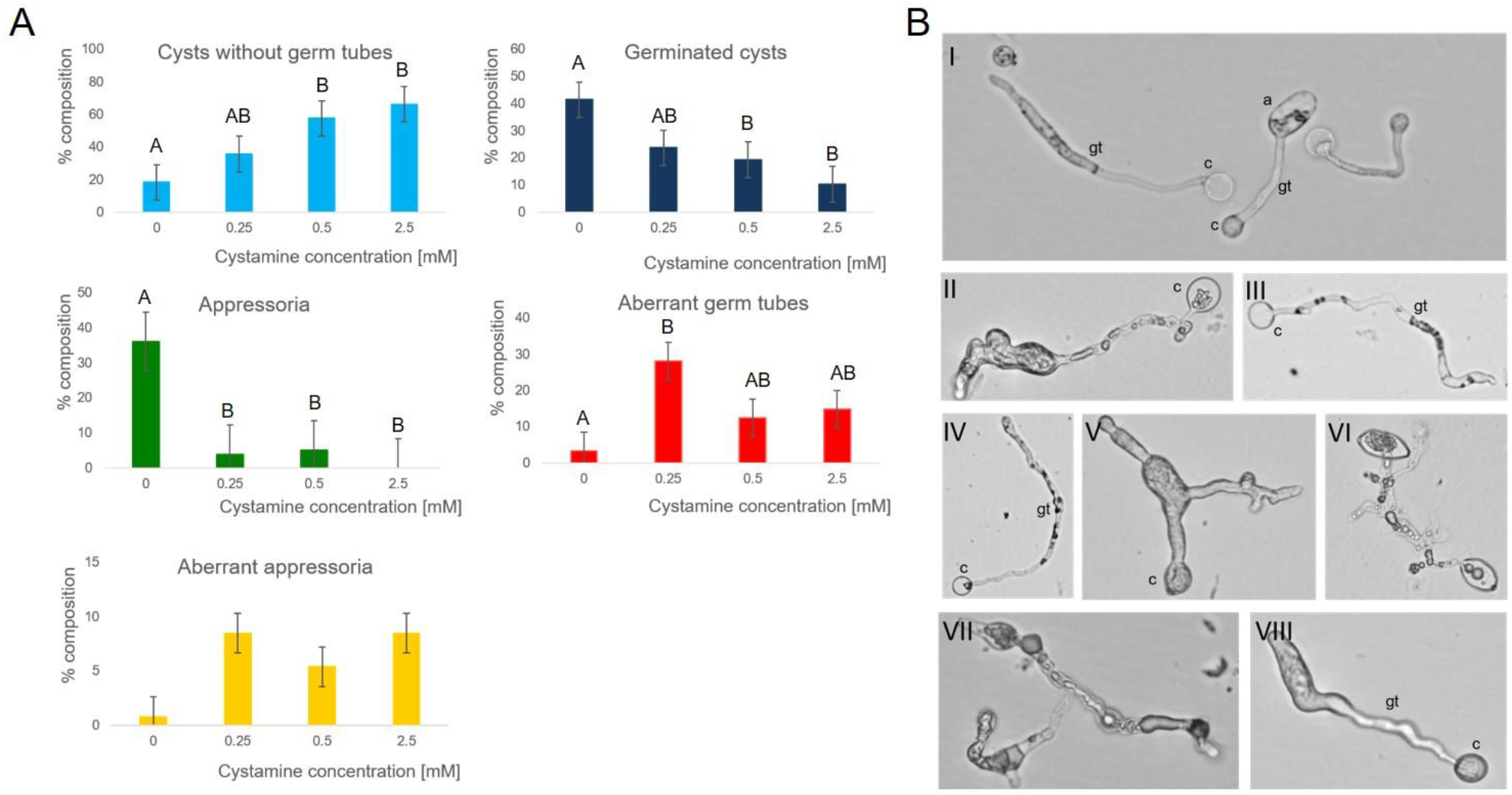
Effect of cystamine treatment on cyst germination and formation of appressoria. **A:** The percentage of cysts, germinated cysts, appressoria, aberrant germ tubes and aberrant appressoria was calculated for samples treated with different concentrations of cystamine. The error bars represent standard error and the letters represent significance of the difference of means (ANOVA followed by Fisher LSD test). **B:** Inverted light microscope images of cysts treated with different concentrations of cystamine and allowed to form appressoria. **I**: wild type, untreated cysts (c) with healthy germ tubes (gt) and appressoria (a); **II:** cysts treated with 0.25 mM cystamine, about 50% germ tubes showed swelling and other deformations; **III and IV:** 0.5 mM cystamine, a large number of cysts germinated, but most of the germ tubes were deformed or collapsed; **V and VI:** 1 mM cystamine, severe deformations to germ tubes, in some cases making it difficult to discern particular structures; **VII and VIII:** 2.5 mM cystamine, severe deformations like in 1 mM cystamine, very few appressoria and the ones that formed were collapsed.

The number of appressoria formed was reduced significantly from about 35 % of all structures in untreated samples to about 5 % in those treated with 0.25 mM and 0.5 mM cystamine. Appressoria formation was severely inhibited by 2.5 mM cystamine (Figure 5A), and the few appressoria that were produced were severely deformed, collapsed or burst (Figure 5B). Cysts treated with cystamine at concentrations higher than 1 mM lost pathogenicity and there were no lesions observed at the site of inoculation on potato leaves under these conditions (Figure 6A and B). Cysts treated with 1 mM and 0.5 mM cystamine produced smaller lesions than the control ones (seen particularly well in Figure 6B) and there was no mycelial growth on the surface of the leaf at the site of inoculation (Figure 6A and B). We verified that the cystamine itself did not have any visual effect on the leaflets and hence that the observed tissue damage was solely due to the pathogen infection. Expression of all six TGase genes was lower in the cystamine treated samples than in the untreated control. The differences in gene expression were not significant between the 0.5 mM and 1.0 mM cystamine treatments (Figure 6C).

**Figure 6.**
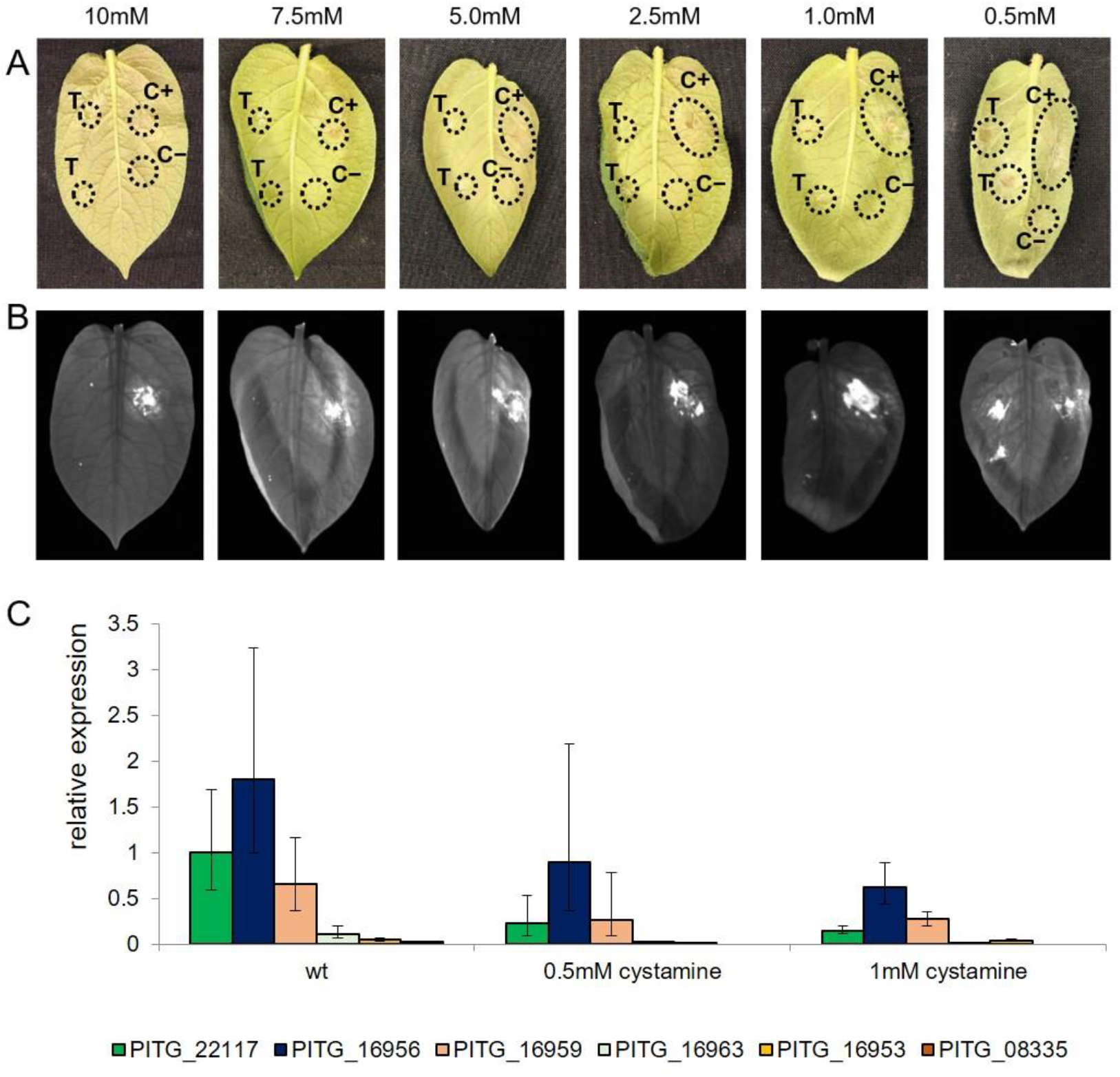
Chemical inhibition of TGases. **A:** Detached leaf assay (DLA). Potato leaflets were inoculated with two drops of cyst solution treated with cystamine of different concentrations (indicated above the pictures) on the left side of the leaf (T), with a control cysts solution in water on the top of the right leaf side (C+) and just cystamine on the bottom right (C-). The same leaf was inoculated with the sample and both controls to screen for possible adverse effects of leaf detachment. Pictures were taken at 3dpi. **B:** DLA leaves scanned on ChemiDoc MP Imager with Cy7 filter (transmission between 755 and 777 nm). Exposure time 5 seconds. The damaged areas of the leaflet appear bright in the image. **C:** Expression of elicitor TGases after treatment with cystamine. Expression of the six putative TGases was tested in samples collected 6 hpi with *P. infestans* cysts treated with cystamine. The expression of the gene was analysed by qRT-PCR with *ActA* as the reference gene and the wt PITG_22117 sample as the calibrator sample set to 1. Error bars represent errors calculated using the modified Delta-Delta Ct method.

Finally, we have also tested the effect of cystamine on protoplast recovery. In line with previous findings in the true fungi (Ruiz-Herrera et al., 1995), treatment with cystamine delayed the recovery of protoplasts. The first emergence of germ tubes was observed after about 36 hours in samples treated with cystamine, whereas in the wild type protoplasts recovery and germination was observed much earlier at about 24 hours (data not shown). This effect was observed in samples treated with 5 mM and 2.5 mM cystamine, but not in lower concentrations of the drug, which corresponds well with the reduction of germination rate and increase in structural abnormalities in sporangia and cysts treated with cystamine.

### P. infestans mycelial extracts exhibit transglutaminase enzymatic activity

Despite sequence and structural differences between the *P. infestans* and human transglutaminases we were able to demonstrate transglutaminase enzymatic activity in crude extracts of *P. infestans* mycelial cultures when tested with a kit designed for human transglutaminases (Abcam Transglutaminase Activity Kit). The enzymatic activity in our samples was lower than would be expected for human tissue samples, yet it was significantly above the detection levels of the kit and at a level similar to the positive control supplied with the kit (Supplementary Figure 1). No significant differences were detected between the cystamine treated (10 mM) and the control samples, supporting the hypothesis that cystamine has only a limited effect on the overall transglutaminase activity of a cell or that it only affects some and not all of the transglutaminases present in the organism.

### Inhibition of TGases reduces turgor pressure of appressoria

Appressoria are structures produced specifically to build pressure and to provide a focal point for the secretion of digestive enzymes to penetrate host cells. To break the barrier of the plant cell wall appressoria must possess a thick cell wall and produce significant turgor pressure, i.e. the pressure resulting from plasma membrane being pushed against cell wall (Wang et al., 2005, Meng et al., 2009). Since we had observed appressoria bursting and cell collapse when *P. infestans* was treated with cystamine *in vitro*, we decided to measure the difference in turgor pressure produced in treated and untreated appressoria, using an incipient plasmolysis cell-collapse assay. The untreated healthy appressoria exhibited an average turgor pressure of 3.325 MPa, while in appressoria produced in the presence of cystamine the turgor pressure was estimated to be 1.327 MPa at 0.25 mM cystamine and 1.282 MPa at 0.50 mM cystamine (Figure 7). This is compared to turgor pressure in the hyphae, which previous studies have estimated to be 0.6-0.8 MPa in fungi and 0.8-1.2 MPa in oomycetes (Brand, 2012, Money, 1990). This significant decrease in turgor pressure of cystamine treated appressoria is in line with our finding that at this concentration pathogenicity of the cysts is largely reduced and clearly demonstrates that an increase in turgor pressure in appressoria is required for infection.

**Figure 7.**
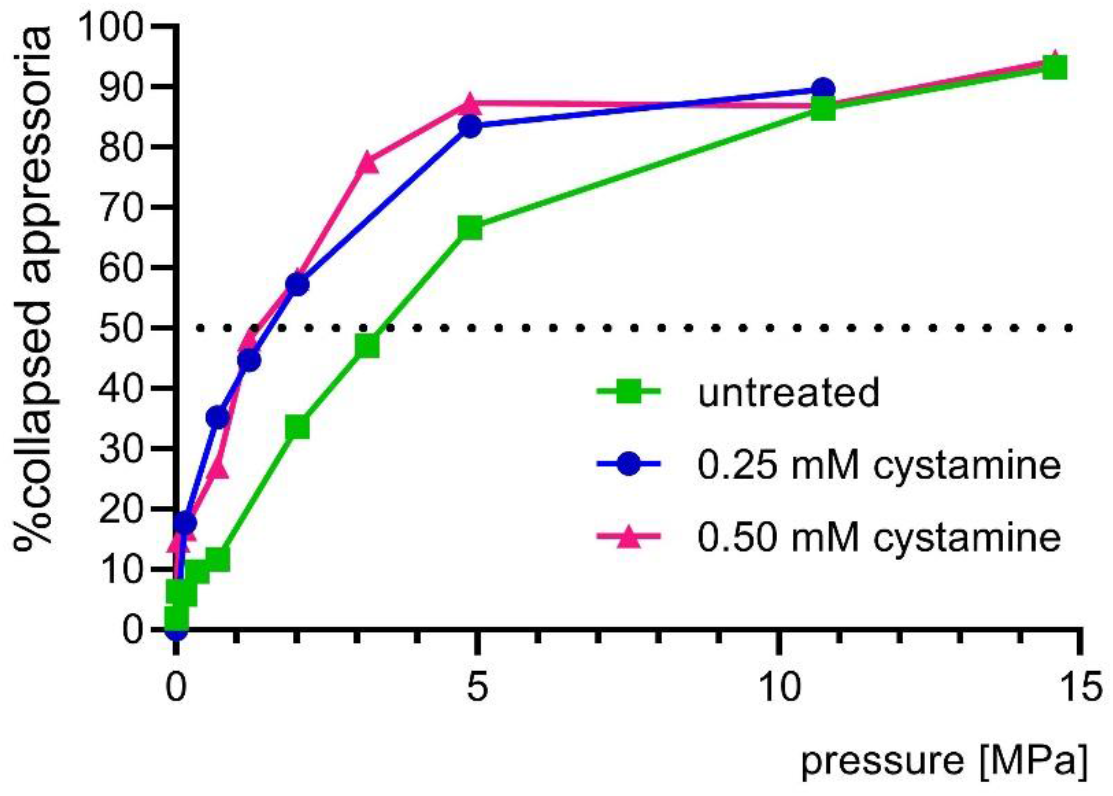
Effect of cystamine treatment on turgor pressure of appressoria. Appressoria were incubated in different concentrations of PEG8000, thus subjected to osmotic pressure ranging from 0 to 15MPa. The turgor pressure of the cell is estimated to be the equal to the pressure at which 50% of appressoria collapse (incipient plasmolysis).

### RNAi-based transient silencing of TGases can be lethal

To confirm the importance of the elicitor-TGases for the viability and development of cysts and appressoria in *P. infestans* we silenced the TGase elicitor genes transiently using RNAi-based protocol. We designed primers to amplify dsRNA molecules that recognise seven of the investigated genes (due to sequence similarity the PITG_16958 gene that showed lack of expression was also targeted). These genes exhibit such high sequence similarity that it was not possible to target them individually with RNAi. Silencing of the entire family of TGases proved lethal and we could not recover the silenced lines. The few lines for which we were able to recover tiny pieces of mycelia grew slowly on a solid medium and the cysts produced from them burst during incubation in water. Given that the high level of sequence conservation prohibits silencing of individual genes, we re-designed the reverse primer to lower its specificity, in an attempt to achieve less-efficient gene silencing. The new reverse primer matched genes PITG_22117 and PITG_16956 100%, but not the other ones. This lower primer specificity allowed us to recover lines with intermediate phenotypes and varying expression levels of the targeted transglutaminase genes (Table 1 and Supplementary Figure 2). The most severe phenotype seen was still lethal with either all of the structures burst or bursting at the time of phenotype assessment (Figure 8B: VI and VII). Several lines displayed phenotypes similar to the control lines with almost all cysts germinating and about 40-50 % of them producing appressoria with few abnormal structures (Figure 8A; lines 2, 9, 12, 15 and 17). The majority of intermediate phenotypes displayed more structural abnormalities than the control lines (Figure 8A; lines 1, 3-8, 13, 14, 16), mostly swelling of the germ tubes (Figure 8B: II and III), but also twisted and very long germ tubes or multiple germ tubes originating from the same cyst (Figure 8B: II). We have also seen structures we previously referred to as “appressoria like” (Grenville-Briggs et al., 2008) – a bulging of the germ tube in an attempt to form an appressorium that fails and the germ tube growth continues (Figure 8B: III-V). Finally, in some of the intermediate phenotypes while there were not many abnormalities observed there were fewer appressoria produced compared to the non-endogenous control lines (Figure 8A; lines 10 and 11). These phenotypes were similar to, and consistent with, those seen by cystamine treatment and described above. All RNAi lines that displayed phenotypic differences compared to the control lines had an overall lower level of transglutaminase gene expression (Table 1 and Supplementary Figure 2). We investigated two lines with higher number of abnormal structures, lines 3 and 7, two lines in which there were less appressoria produced but not many abnormalities, lines 10 and 11, and two lines with a wild type phenotype, lines 9 and 12. As predicted from the expression profiles in life cycle stages and infection time points (Figure 3) the PITG_16958 gene was not expressed in any of the samples. In line 3 all the remaining six genes were expressed at lower level compared to the control and in line 7 four genes showed lower expression and two showed expression levels similar to, or higher than, the control lines (Table 1 and Supplementary Figure 2). In the lines with fewer appressoria, lines 10 and 11, the overall expression was also lower than in the control sample, with only one gene in line 11 and two in line 10 showing similar or slightly higher expression levels than the controls (Table 1 and Supplementary Figure 2). In the two lines that did not display phenotypic differences the overall expression of transglutaminases was higher than in the other RNAi lines, with three genes showing expression similar to, or higher than control levels, and three genes showing reduced expression in line 9; while in line 12 five out of six genes were expressed at control levels or higher (Table 1 and Supplementary Figure 2).

**Table 1.** Relative expression of transglutaminase genes in RNAi lines. The presented values are expression relative to the expression of the reference *Actin A* gene and calibrated to the expression of the specific gene in the GFP control sample, which was set to 1. Values above 1 (pink) indicate higher and below 1 (purple) lower expression than the expression of the given gene in the control sample. * indicates lack of detected signal, i.e. the expression is below the detection level. Pathogenicity data from DLA experiments are presented for comparison.

| RNAi line | Gene number |  |  |  |  |  | Pathogenicity |
| --- | --- | --- | --- | --- | --- | --- | --- |
|  | PITG_22117 | PITG_16956 | PITG_08335 | PITG_16953 | PITG_16959 | PITG_16963 |  |
| **3** | * | * | * | * | * | * | Lower |
| **7** | 1.945746 | 0.303926 | 0.507534 | 1.964669 | 0.301265 | 0.324826 | Lower |
| **9** | * | 1.98087 | * | 1.779605 | 2.041624 | * | Yes |
| **10** | 0.892556 | 0.33275 | 0.604045 | 1.668856 | 1.103508 | * | Lower |
| **11** | 1.057386 | 0.565048 | * | 0.929691 | 0.698291 | 0.295875 | Lower |
| **12** | 5.911655 | 2.406242 | 2.300192 | 0.970109 | 2.925382 | 1.577009 | Yes |

**Figure 8.**
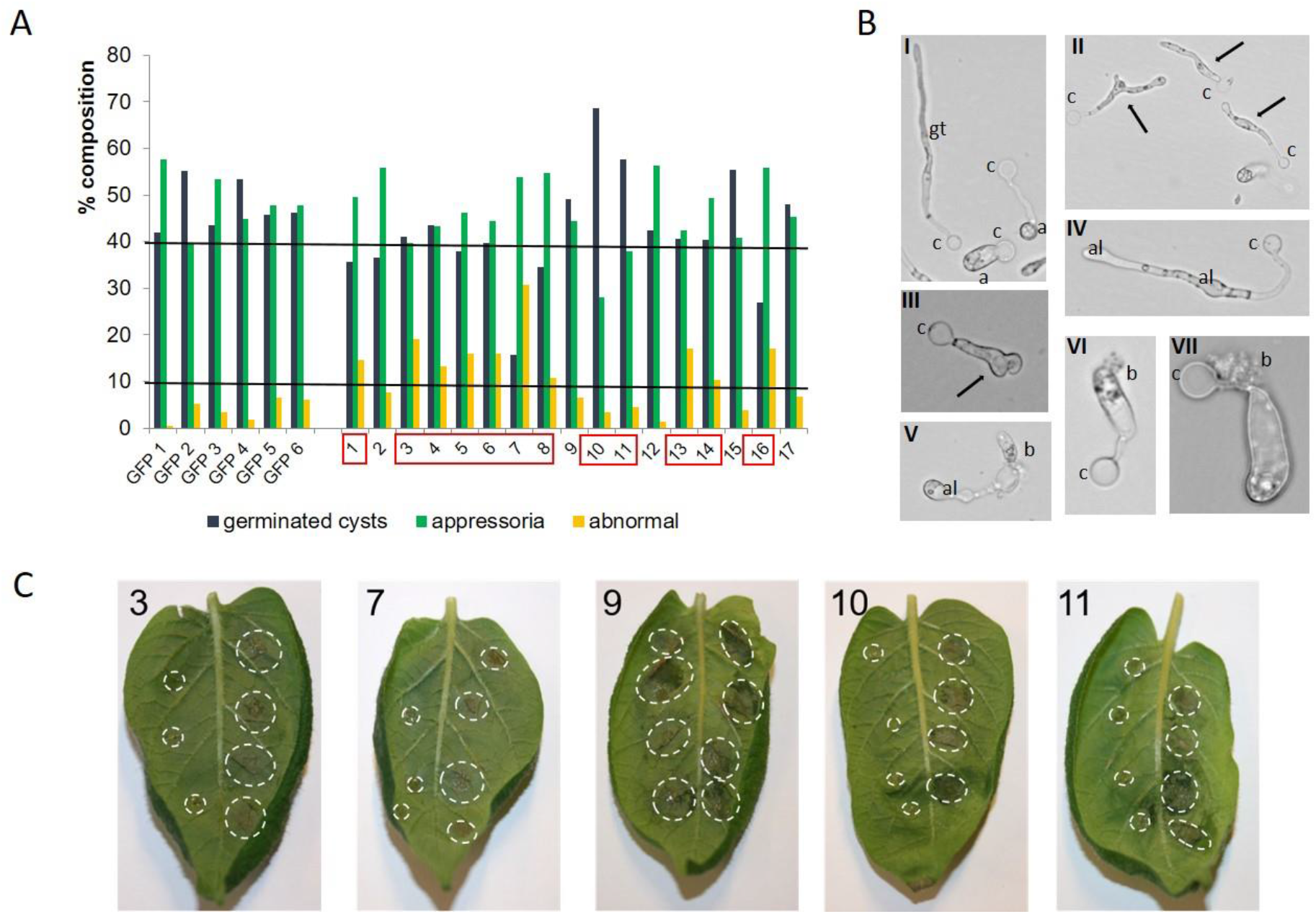
RNAi-based transient silencing of elicitor-TGases. **A:** Percentage of germinated cysts, appressoria and abnormal structures counted in individual silenced lines and individual GFP controls. The lines marked in red were less pathogenic than the control ones. The horizontal lines are arbitrary thresholds for the minimal number of healthy appressoria (upper) and maximum number of abnormal structures (lower) for a line to be pathogenic. **B:** Inverted light microscope images of the counted structures. I: GFP controls with healthy cysts (c), germ tubes (gt) and appressoria (a); II and III: swelling and multiple germ tubes; IV and V: apressoria-like structures (al); V, VI and VII: bursting of appressoria and cyst **C:** Detached leaf assay (DLA). Potato leaves were inoculated with four drops of cyst solution from different RNAi lines on the left side and GFP control lines on the right side of the main vein. The RNAi line is indicated in the top left corner of each picture. Pictures were taken at 4dpi.

To test the pathogenicity of the RNAi lines, cysts were collected from the same lines that were used for phenotypic assessment. For detached leaf assays the same leaf was inoculated with cysts from a single TGase RNAi line on the left side and from a single GFP non-endogenous control line on the right side of the primary vein to account for the possible adverse effects of the leaf detachment, or minor variations in individual leaves. The majority of the TGase RNAi lines showed reduced or no pathogenicity when compared to the control lines, including lines 3, 7, 10 and 11 (Figure 8C) in which expression of transglutaminase genes was overall lower than in the controls (Table 1 and Supplementary Figure 2). The pathogenicity of lines 12 and 19 was not affected, which corresponds well with the gene expression data (Table 1 and Supplementary Figure 2). Comparison of the phenotype, pathogenicity and gene expression data suggests that neither of the investigated TGases is essential for the development of cysts and appressoria on its own, but a certain expression level is necessary, i.e. the proteins are redundant to some extent. Nonetheless, decreased expression of the PITG_16956 gene in all of our RNAi lines with decreased pathogenicity points towards the role of this particular gene in appressoria development and is consistent with its high expression in germinated cysts (Figure 3). The similarity of the phenotype observed in the silenced lines and the cysts treated with cystamine validates the hypothesis that whilst complete inhibition of TGase activity is lethal to *P. infestans*, the redundancy of the proteins allows for partial loss of function in any of them.

### Conclusions

The presented data shows that transglutaminases are essential for the development of a healthy cell wall and thus growth, development and pathogenicity of *P. infestans*. The chemical inhibition and RNAi silencing assays proved that targeting of these TGases could be used as an efficient control method. Targeting transglutaminase function to control late blight disease may offer a highly specific and potentially durable method of disease control. However, to succeed, all of the transglutaminase genes would need to be targeted simultaneously to avoid the potential risk of *P. infestans* overcoming such a pesticide by adaptation of one of these partially redundant genes. Our data also show RNAi-based silencing to be a powerful method for the evaluation of potentially essential genes for which stable transformation and full silencing would prove lethal. Finally, for the first time we have shown that *P. infestans* appressoria build up turgor pressure, which although not at the level of that seen in fungal phytopathogens, appears nevertheless to be essential to the infection process.

## Experimental Procedures

### In silico analysis and phylogenetics

The DNA sequence of *Phytophthora infestans* gene PITG_22117 was used to search for similar sequences in *P. infestans* using BLASTn. Alignment of the sequences was performed using Clustal Omega (Madeira et al., 2019). To identify all proteins predicted to have TGase function string search “*Phytophthora infestans* transglutaminase” was used to search the NCBI Gene database. For all 21 results the protein sequences were retrieved from the database and used to construct a maximum likelihood phylogenetic tree (bootstrap method, 1000 replications) using MEGA-X software (Kumar et al., 2018). The functional domain and signal peptide predictions were performed using InterProScan (Jones et al., 2014).

### Phytophthora infestans cultivation

All experiments were carried out using *P. infestans* strain 88069. For maintenance, cultures were grown on solid rye sucrose medium (Caten and Jinks, 1968), at 18 °C in darkness and sub-cultured every two to three weeks. Liquid cultures for extraction of RNA and growth inhibition assays were grown in pea broth, pH 7.25, at 18 °C in darkness.

### Transglutaminase expression throughout P. infestans life cycles and infection time course

To analyse the expression profiles of all the investigated transglutaminases, 14-day old cultures were grown on rye sucrose and sporangia, zoospores, cysts, germinated cysts and appressoria were collected as described previously (Grenville-Briggs et al., 2010, Resjö et al., 2017). The samples were collected from pooled material originating from 20 individual cultures. The collected cysts were used to inoculate potato leaves (cultivar Désirée) in a detached leaf assay and infection time point samples were collected at: 6 hpi, 12 hpi and 24 hpi. Mycelium samples were grown in liquid pea medium for 48 h before collection. Each of the samples was ground in liquid nitrogen and the total genomic RNA was extracted using the Qiagen RNeasy Plant Mini kit following the manufacturer’s protocol. The samples were DNase treated (Turbo DNA-free Kit, Invitrogen) before first strand cDNA synthesis was performed as described (SuperScript III, Invitrogen; Grenville-Briggs et al., 2010). All cDNA samples were diluted to 5 ng μl^-1^ and the infection samples to 20 ng μl^-1^ before gene expression analysis by qRT-PCR. The qRT-PCR was performed in a BioRad real-time PCR cycler, using SYBR green as the fluorescent dye and the primer pairs listed in Table 2. The expression of transglutaminase genes was normalised to a reference gene *Actin A* and mycelium was used as a calibrator sample with expression set to 1, the relative expression was calculated using modified Delta-Delta Ct method, as described previously (Avrova et al., 2003).

**Table 2.**
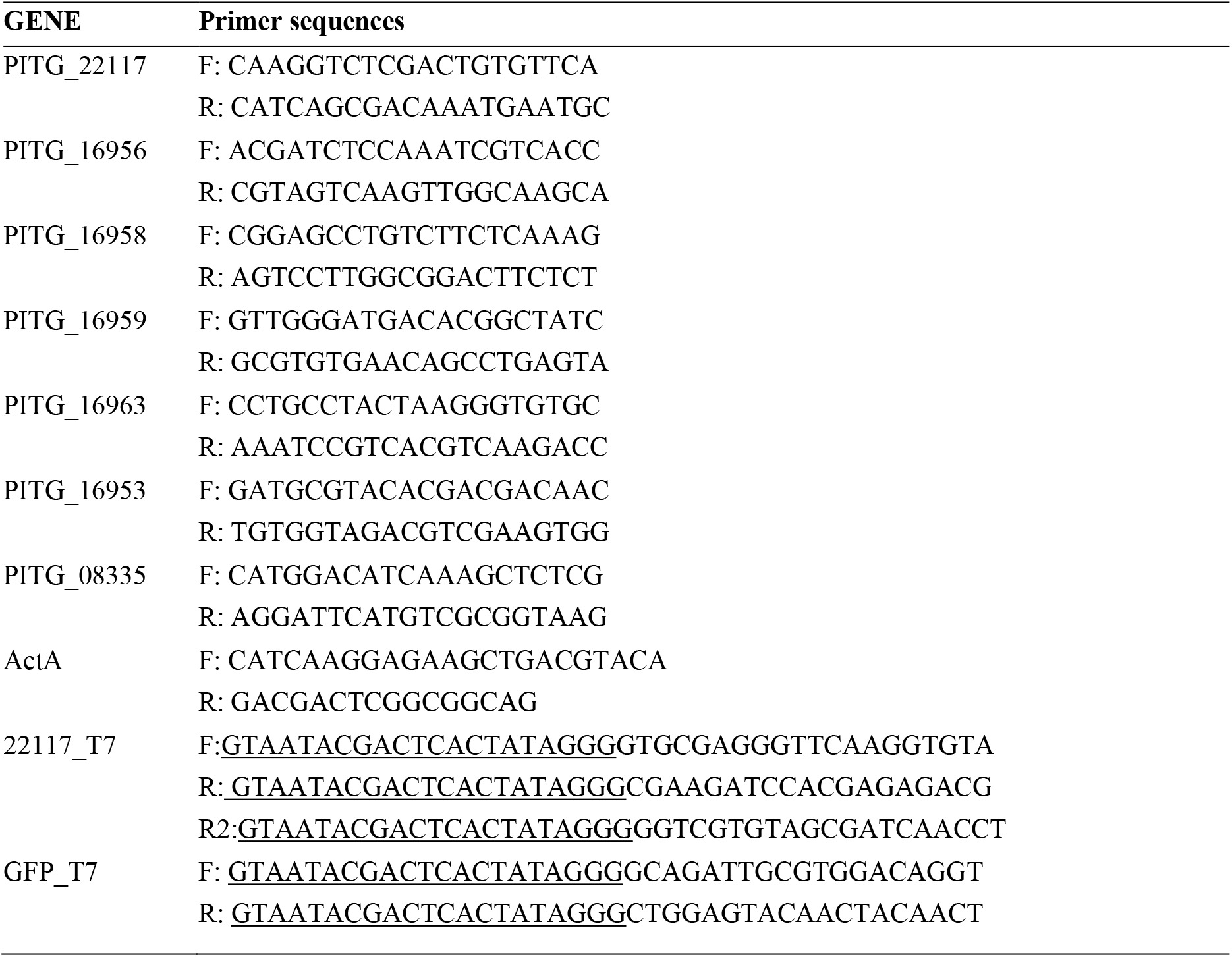
Primer pair sequences used for qRT-PCR and dsRNA synthesis. The underlined fragments of the dsRNA synthesis primers are the T7 promoter sequences.

### RNA interference

Oligonucleotide primers with T7 polymerase RNA promoters (Table 2) were designed to amplify a 200 bp long transglutaminase amplicon. Due to high sequence similarity, it was not possible to design unique primers for each of the transglutaminase genes; therefore, the designed primers were able to amplify fragments from seven similar transglutaminase genes: PITG_22117, PITG_16953, PITG_16956, PITG_16958, PITG_16959, PITG_16963 and PITG_08335. Lack of binding elsewhere was confirmed by BLASTn search within *P. infestans* genome (performed with the low-complexity filter turned off). 1 μg of the PCR product was used to synthesise dsRNA with MEGAscript RNA interference kit (Ambion) according to manufacturer’s protocol. GFP was used as a non-endogenous positive control to ensure that any possible phenotypical changes arise from the silencing and not the transformation protocol. Preparation of protoplasts and introduction of the dsRNA were performed as described (Grenville-Briggs et al., 2008). Fourteen days after the transfection, zoospores were collected from single colony plates and encysted. A small portion (about 200 μl) of each of the samples was used for detached leaf assays and gene expression analysis, whilst the remainder was incubated in plastic petri dishes at 11 °C in darkness for 16 h to induce cyst germination and appressoria formation. The number of germinated cysts, appressoria, and any aberrant structures were counted using an inverted light microscope. The silenced lines were compared to the GFP control lines.

### Assessment of pathogenicity of the RNAi lines

Pathogenicity of the RNAi lines was assessed by detached leaf assays. Healthy leaves of similar size were removed from the middle of the potato plant and placed abaxial side up in plastic boxes lined with moist paper tissue to ensure high humidity levels were maintained. Each leaf was inoculated with four 10 μl droplets of cyst solution from a single RNAi line on one side of the leaf and four 10 μl drops of the GFP-control cysts on the other side of the leaf. There were four leaves inoculated for each RNAi line, two for visual assessment of pathogenicity and two for material collection for gene expression analysis (see next section). The boxes were sealed with parafilm and placed in climate chambers with a cycle of 16 h light and 8 h darkness at 18 °C, starting with the dark period as described in Resjö et al. (2017). The symptoms were compared at 4 dpi and 7 dpi. Visual observations were complemented by leaf scans performed in Bio-Rad ChemiDoc MP Imager (Zahid et al., 2021).

### Transglutaminase expression in RNAi lines

To ensure sufficient amount of material for the gene expression analysis cysts produced from the RNAi lines were used to inoculate potato leaves (as described in the previous section). Eight leaf discs per sample were collected at 6 hpi and snap frozen in liquid nitrogen. The cork borer and forceps used for sample collection were washed with 70% ethanol between different lines. The RNA extraction and qRT-PCR analyses were performed as described above, using primers listed in Table 2.

### Effect of transglutaminase inhibitor cystamine on P. infestans growth and pathogenicity

To test the effect of cystamine on *P. infestans* growth, liquid medium with varying concentrations of cystamine ranging from 0.5 mM to 250 mM, was inoculated with small plugs of solid agar culture that were excised with a cork borer to ensure the same size for all cultures. The growth differences were estimated daily until the control culture reached the edges of the petri dish. To test if the effects of the drug were reversible the cultures containing cystamine were left in the incubator for 14 days. Additionally, a range of cystamine concentrations were added to solid rye sucrose medium and the cultures were grown until radial growth of the control cultures reached the edge of the petri dish. Solid cultures with cystamine in the medium were then used to test the effect of the drug on sporulation. The plates were flooded with cold sterilised tap water and sporangia were collected and counted using a haemocytometer. To investigate the effects of cystamine on zoospore release and motility, 12-14-day old cultures were flooded with either water or cystamine solutions at varying concentrations, and incubated at 4 °C for 4 h, after which zoospores were harvested, filtered through a 40 μm mesh and counted. Encystment was induced as described previously (Resjö et al., 2017) and samples were incubated for 2-4 h at r.t. after which cyst germination was evaluated by light microscopy. Alternatively, to assess the rate of cyst germination, 12-14-day old cultures were flooded with water, incubated at 4 °C to release zoospores, filtered and encysted. The cysts were collected by centrifugation (1200 xg, 15 min), the supernatant was removed and cysts were re-suspended in either fresh water (controls) or cystamine solution. Germination was assessed as described above, after 2-4 h incubation at r.t.

Finally, in order to assess appressorium formation, cysts were induced as described above, either in the presence of cystamine or treated with cystamine after encystment, incubated for 16 h at 11 °C in petri dishes. Control samples were encysted as described and then treated with water. The number of cysts, germinated cysts, appressoria and any aberrant structures were counted using an inverted light microscope.

The effect of the cystamine treatment on *P. infestans* pathogenicity was tested with DLA as described above for RNAi. The leaves were inoculated with two droplets each containing 50 000 cystamine-treated cysts on the left of the central vein and on the right side with a water-treated control cysts and additionally with just cystamine to test the effect of the drug on the potato leaf.

### TGase gene expression in cystamine-treated cysts

Leaf disc samples were collected at the inoculation site of the DLA assay (described above) at 6hpi. Each sample consisted of eight leaf discs collected from three leaves. The RNA extraction, DNase treatment, cDNA synthesis and qRT-PCR were performed as described above.

### Effect of cystamine on appressorium turgor pressure

Turgor pressure of the appressoria was measured indirectly by counting the number of plasmolysed appressoria in various concentrations of PEG8000 using an incipient plasmolysis assay (Howard et al., 1991, Michel, 1983). These concentrations covered a range of osmotic pressures, which were represented using a standard curve. The turgor pressure of the cell is estimated to be the equal to the pressure at which 50 % of appressoria collapse (incipient plasmolysis). The graph and calculations were done in GraphPad Prism 8.2.1, using the standard curve interpolation. Separate curves were drawn for samples with and without cystamine.

### Transglutaminase enzymatic activity assay

*P. infestans* mycelial cultures were grown in liquid pea broth medium in the presence of 10 mM cystamine and without cystamine addition (control). 3-day old cultures were blotted on sterile filter paper and ground using plastic micropestles and sterile sand. The crude extracts were assessed for the transglutaminase enzymatic activity using Abcam Transglutaminase Activity Kit (ab204700) according to manufacturer’s protocol.

## Supporting information

Supplementary Figure 1. Transglutaminase activity in P. infestans. The transglutaminase activity was measured using Abcam kit (ab204700) where the enz

Supplementary Figure 2. Expression profiles of the elicitor-Tgases in RNAi-silenced lines. The expression was calculated relative to ActA and calibrat

## Acknowledgements

This project has received funding from the European Union’s Horizon 2020 research and innovation programme under Grant Agreements No 774340 (Organic Plus) and No 766048 (MSCA-ITN-2017 PROTECTA), as well as the the Swedish Research Council FORMAS (grant 2019-00881 to LGB). The authors would also like to thank Hadis Mostafanezhad for her help with microscopic observations and Kristian Persson Hodén for his input in the discussion on transglutaminase expression in *P. infestans* oospores.

