## Supplementary figures and images for "A family of cell wall transglutaminases is essential for appressorium development and pathogenicity in *Phytophthora infestans*"

### Supplementary Figure 1. Transglutaminase activity in P. infestans. The transglutaminase activity was measured using Abcam kit (ab204700) where the enz

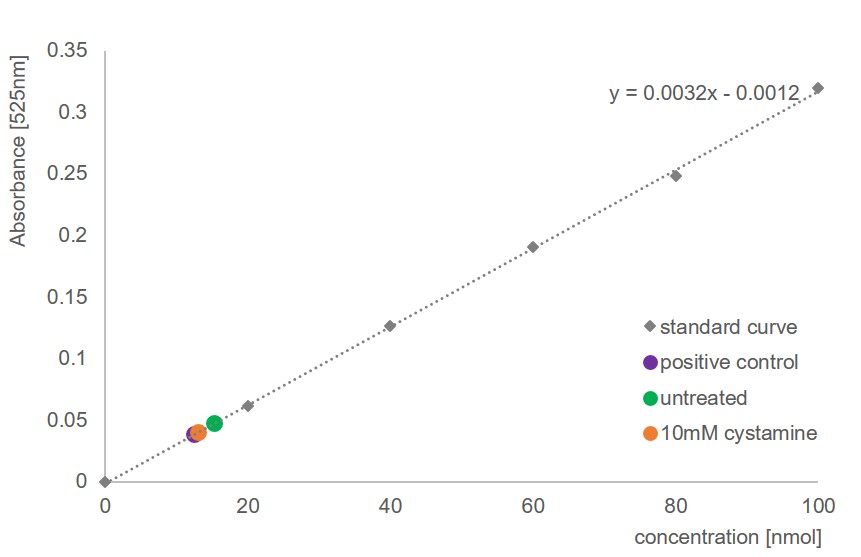

### Supplementary Figure 2. Expression profiles of the elicitor-Tgases in RNAi-silenced lines. The expression was calculated relative to ActA and calibrat

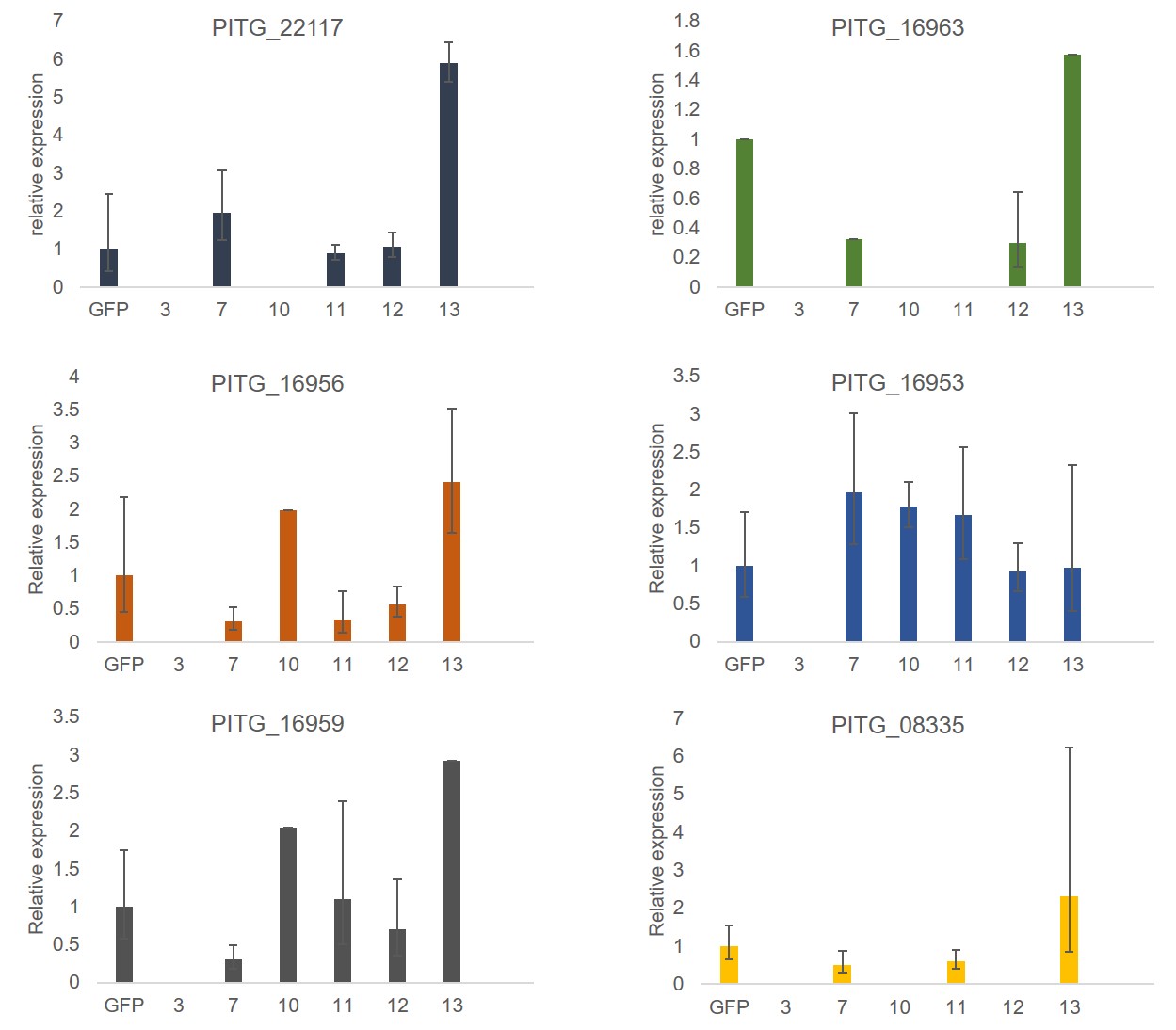
